# Multiple reference genome sequences of hot pepper reveal the massive evolution of plant disease resistance genes by retroduplication

**DOI:** 10.1101/115410

**Authors:** Seungill Kim, Jieun Park, Seon-In Yeom, Yong-Min Kim, Eunyoung Seo, Ki-Tae Kim, Myung-Shin Kim, Je Min Lee, Kyeongchae Cheong, Ho-Sub Shin, Saet-Byul Kim, Koeun Han, Jundae Lee, Minkyu Park, Hyun-Ah Lee, Hye-Young Lee, Young-sill Lee, Soohyun Oh, Joo Hyun Lee, Eunhye Choi, Eunbi Choi, So Eui Lee, Jongbum Jeon, Hyunbin Kim, Gobong Choi, Hyeunjeong Song, JunKi Lee, Sang-Choon Lee, Jin-Kyung Kwon, Hea-Young Lee, Namjin Koo, Yunji Hong, Ryan W. Kim, Won-Hee Kang, Jin Hoe Huh, Byoung-Cheorl Kang, Tae-Jin Yang, Yong-Hwan Lee, Jeffrey L. Bennetzen, Doil Choi

## Abstract

Transposable elements (TEs) provide major evolutionary forces leading to new genome structure and species diversification. However, the role of TEs in the expansion of disease resistance gene families has been unexplored in plants. Here, we report high-quality *de novo* genomes for two peppers (*Capsicum baccatum* and *C. chinense*) and an improved reference genome (*C. annuum*). Dynamic genome rearrangements involving translocations among chromosome 3, 5 and 9 were detected in comparison between *C. baccatum* and the two other peppers. The amplification of *athila* LTR-retrotransposons, members of the *gypsy* superfamily, led to genome expansion in *C. baccatum.* In-depth genome-wide comparison of genes and repeats unveiled that the copy numbers of NLRs were greatly increased by LTR-retrotransposon-mediated retroduplication. Moreover, retroduplicated NLRs exhibited great abundance across the angiosperms, with most cases lineage-specific and thus recent events. Our study revealed that retroduplication has played key roles in the emergence of new disease-resistance genes in plants.

## Introduction

Long terminal repeat retrotransposons (LTR-Rs) are a major evolutionary force in animals, fungi, and, especially, plants. They comprise >75% of many plant genomes and cause genomic instability, including genome expansion by amplification using an RNA intermediate^1^. Besides genome expansion, LTR-Rs facilitate the creation of new candidate genes called retrogenes by means of retroduplication, in which spliced mRNA is captured, reverse transcribed, and subsequently integrated into the genome by a retrotransposon^2-4^. In contrast to transduplication caused by class II transposable elements (TEs)^5,6^, the distinctive features of retrogenes are (i) intron loss compared to their parental source genes, (ii) the presence of a 3’ poly(A) tail, and (iii) flanking direct repeats^7^.

The evolutionary forces acting on most plant retrogenes are still largely unclear^3, 8-11^. Although LTR-Rs are the most abundant TEs in all but the tiniest plant genomes, few studies have been reported on the detection of retrogenes generated by LTR-Rs in plants^6,12,13^. Wang *et al*^3^ identified 27 retrogenes within LTR-Rs in rice and concluded that retrogenes that originated within LTR-Rs were often not classified as retrogenes, partly because of the rapid destruction of the LTR-R structure by illegitimate recombination^14^. Moreover, they suggested that the retrogenes might be very frequent in grass species due to the abundance of LTR-Rs. In agreement with this prediction, recent studies have reported the genome-wide identification of hundreds retrogenes within LTR-Rs in maize^15^, rice and sorghum^16^ as well as the existence of retrogenes captured by LTR-Rs in *Arabidopsis*^4^. However, most of those retrogenes were classified as pseudogenes or uncharacterized genes.

Previous studies reported the massive capture of specific gene families by certain TEs and suggested a correlation between TE-mediated gene duplication and specific gene family expansion^17,18^. The nucleotide-binding and leucine-rich-repeat proteins (NLRs) represent a highly amplified gene family in plants and provide the majority of functional plant disease resistance loci^19-21^. Comparative genomic analyses have suggested the possibility of LTR-Rs and NLRs co-evolution, partly because they are often co-localized^20,22,23^. Because the NLRs usually reside in clusters within genomes, NLR expansions have been mainly interpreted as the products of ectopic recombinational duplications^19,24^.

Here, we report high-quality *de novo* genomes of two novel domesticated *Capsicum* species and also improved the quality of the reference pepper genome^25^. Comparative analyses of the three pepper genomes, with other plant genomes as outgroups, provided insights into genome evolution and species diversification in the genus *Capsicum.* Our analyses unveil an important mechanism for the massive emergence of new plant NLRs by LTR-R-mediated retroduplication and show the dynamic evolutionary processes for functional disease resistance genes in plants.

## Results

### *De novo* sequencing, assembly and annotation of *Capsicum* genomes

We sequenced and assembled the genomes of *Capsicum baccatum* PBC81 (hereafter, Baccatum) and *C. chinense* PI159236 (hereafter, Chinense) using Illumina HiSeq 2500 with library insert sizes ranging from 200 bp to 10 kb (Supplementary Tables 1-3). The estimated genome sizes of Baccatum and Chinense, based on 19-mer analysis, were 3.9 and 3.2 Gb, respectively (Extended Data Fig. 1). The assembled genomes of Baccatum and Chinense constituted 3.2 and 3.0 Gb (83 and 94% of the estimated genome sizes, respectively) and had scaffold (contig) N50 sizes of 2.0 Mb (39 kb) and 3.2 Mb (49 kb), respectively (Supplementary Table 3). We annotated the protein-coding genes in the Baccatum and Chinense assemblies as well as those in the pre-existing *C. annuum* CM334 genome^25^ (hereafter, Annuum) for detailed comparative analysis (Extended Data Fig. 2). On average, ~35,000 genes were annotated in each species (Supplementary Table 4). A comparison of the updated and previous gene models of Annuum revealed ~ 10,000 genes that did not overlap between the two models, suggesting that most of the non-overlapping genes in the previous version were associated with TEs (Extended Data Fig. 3).

**Extended Data Figure 1.**
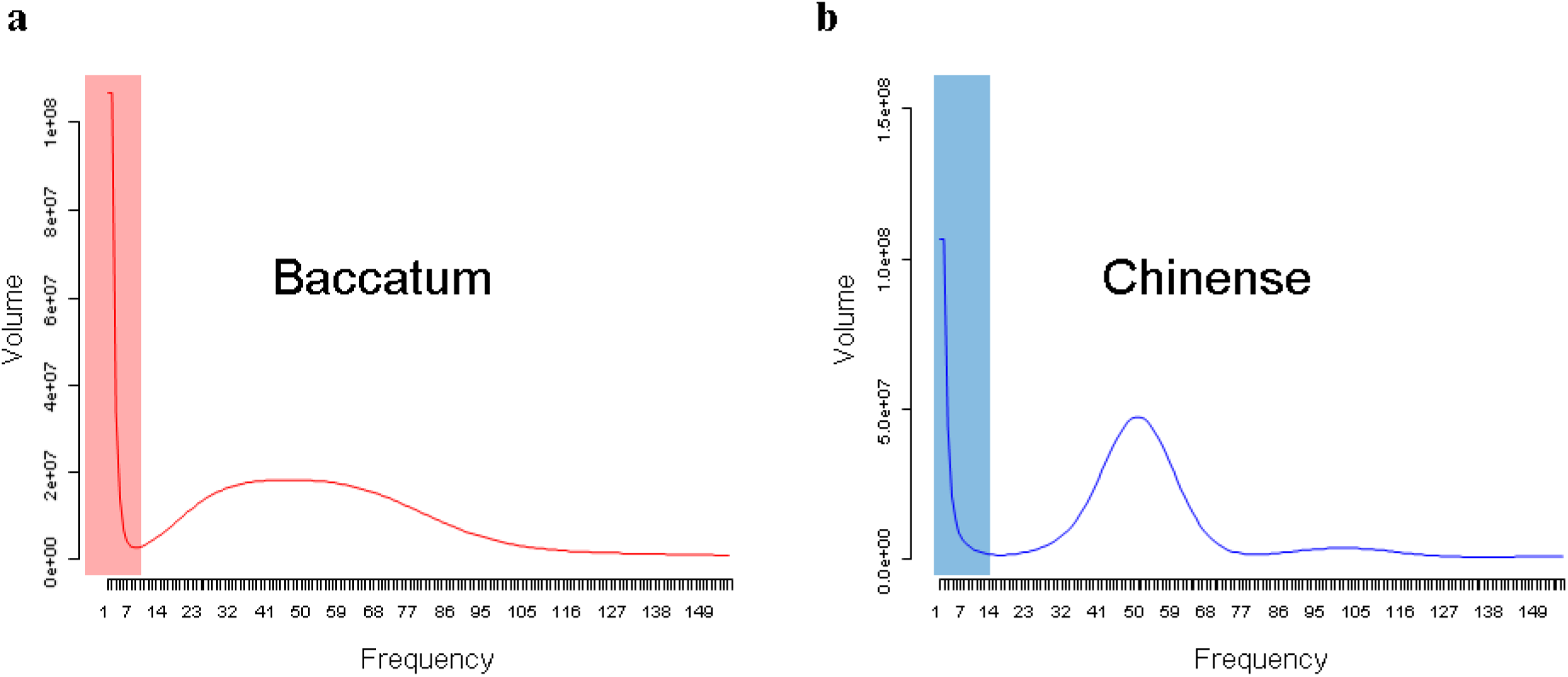
The 19-mer distribution patterns for the Baccatum and Chinense genome sequences. **a-b,** The *x-* and *y-axes* indicate the frequency and volume of 19-mers, respectively. Shaded regions in the graphs indicate low-frequency data as erroneous candidates.

**Extended Data Figure 2.**
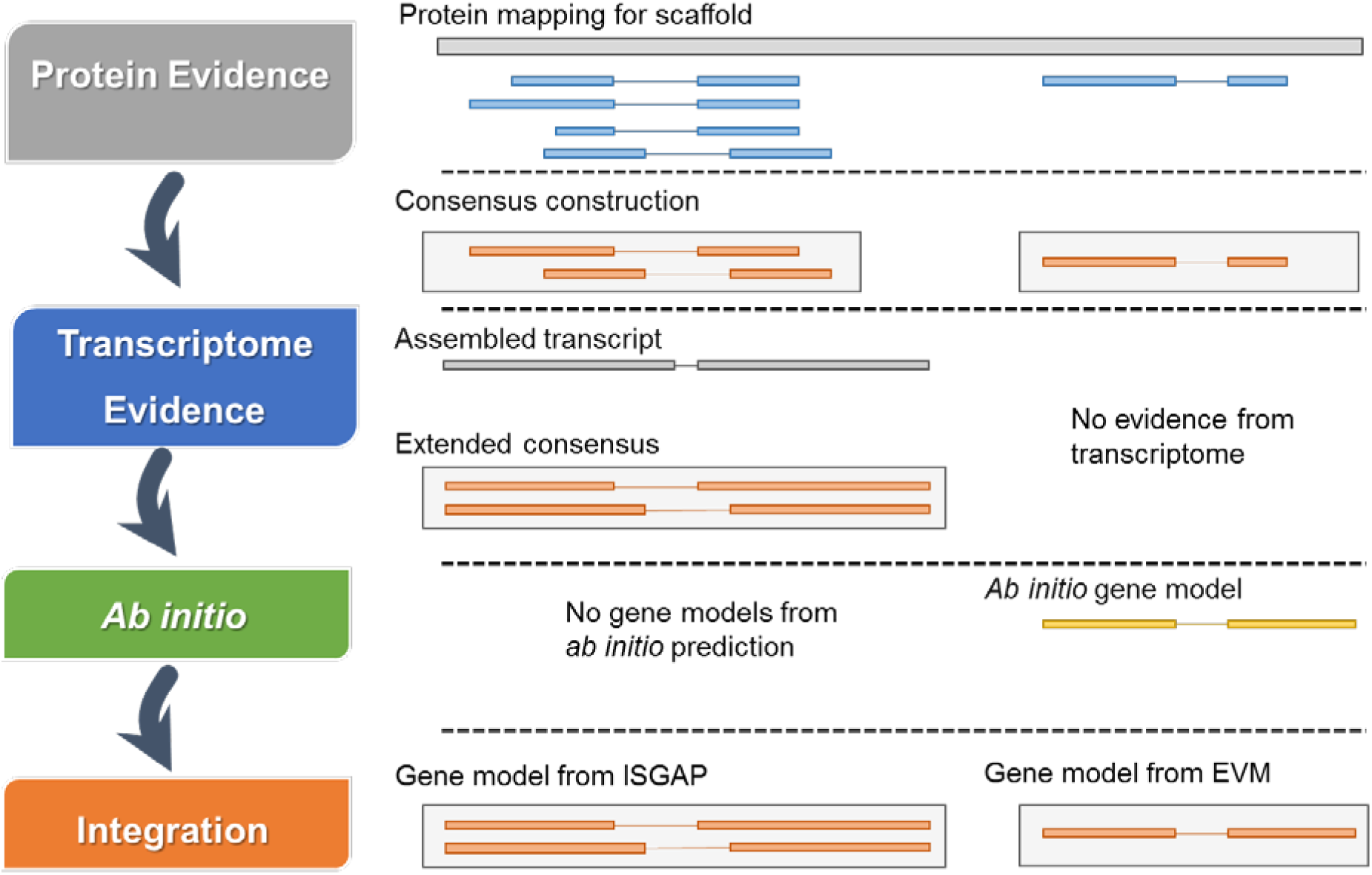
Gene annotation scheme for the pepper genomes. The diagram depicts our gene annotation process for the pepper genomes using proteins, transcriptome and *ab initio* prediction. ISGAP^44^ identifies gene structure based on transcriptome and protein evidences. EVM^48^ integrates results of protein mapping and / or *ab initio* prediction using AUGUSTUS^47^.

**Extended Data Figure 3.**
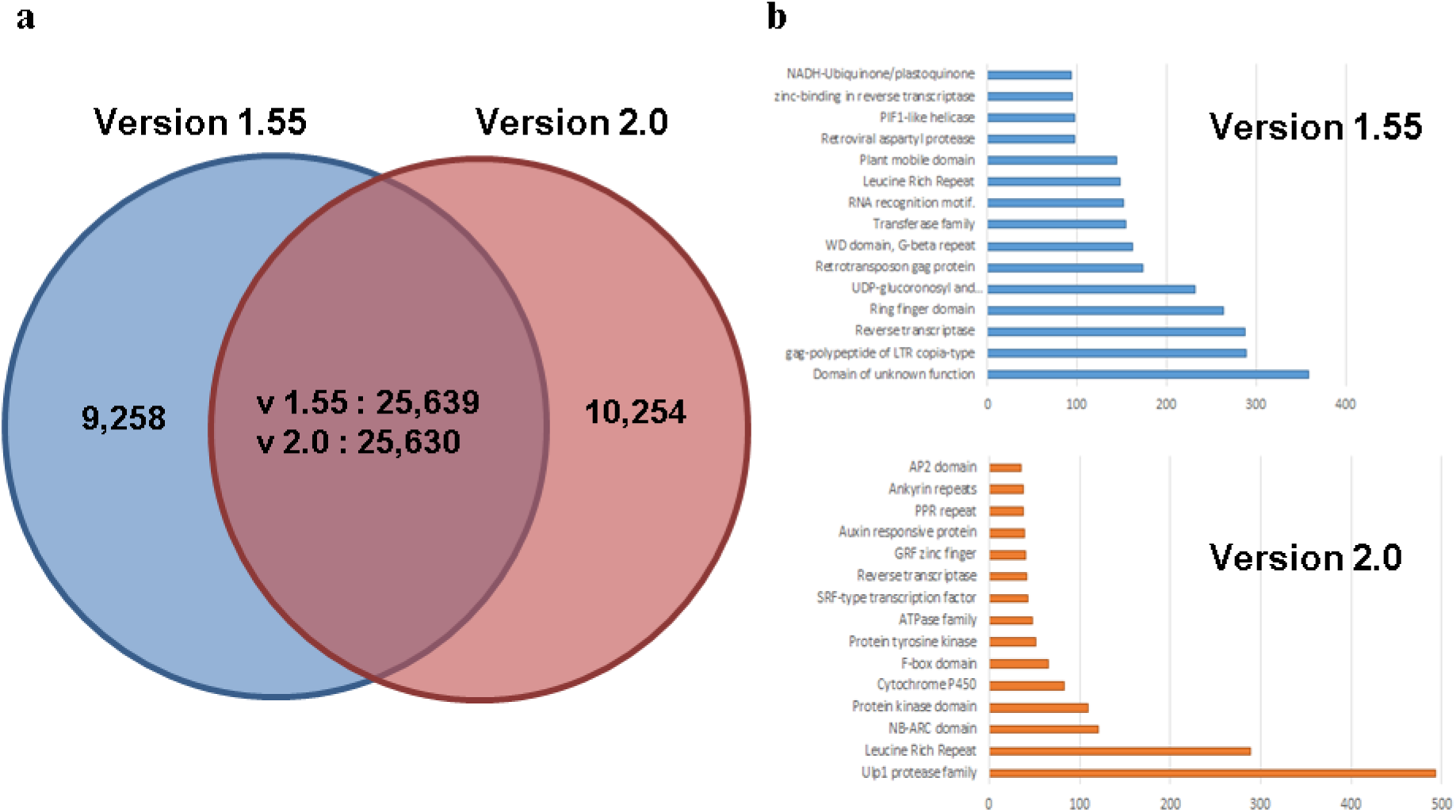
Comparison of annotated gene sets between the pre-existing and newly generated versions of the Annuum genome. **a,** The Venn diagram indicates the numbers of overlapping and non-overlapping gene models between versions 1.55 and 2.0, using the genomic positions as one consideration. **b,** The bar graphs show the numbers of predicted genes in the top 15 domain descriptions in non-overlapping gene models in v1.55 and v2.0. The *x-* and *y-axes* indicate the number of annotated genes with a particular domain and the category of domain, respectively.

A high-density genetic map of each species was generated by genotyping-by-sequencing on F2 populations^26,27^ (Extended Data Fig. 4). After breaking up chimeric scaffolds on the basis of genetic map information, we organized the assembled genomes into 12 pseudo-chromosomes. Overall, 87% of Baccatum (2.8 Gb in 2,083 scaffolds) and 89% of Chinense (2.8 Gb in 1,557 scaffolds) in assembled genomes were ordered by the genetic map and inspected for syntenic inferences with the improved pseudomolecules of Annuum (Supplementary Table 5). We validated the assembled genomes by reference guided mapping using the refined single-end and paired-end data, and alignment to the assembled transcriptome of each species. In total, we detected more than 98.1% of the filtered raw sequences (>98% identity) and more than 93.4% of the assembled transcriptomes (>98% identity and 80% of query coverage) in the genome assemblies (Supplementary Table 6). Taken together, our analyses provide the *de novo* reference genomes of two new pepper species as well as an improved Annuum genome.

**Extended Data Figure 4.**
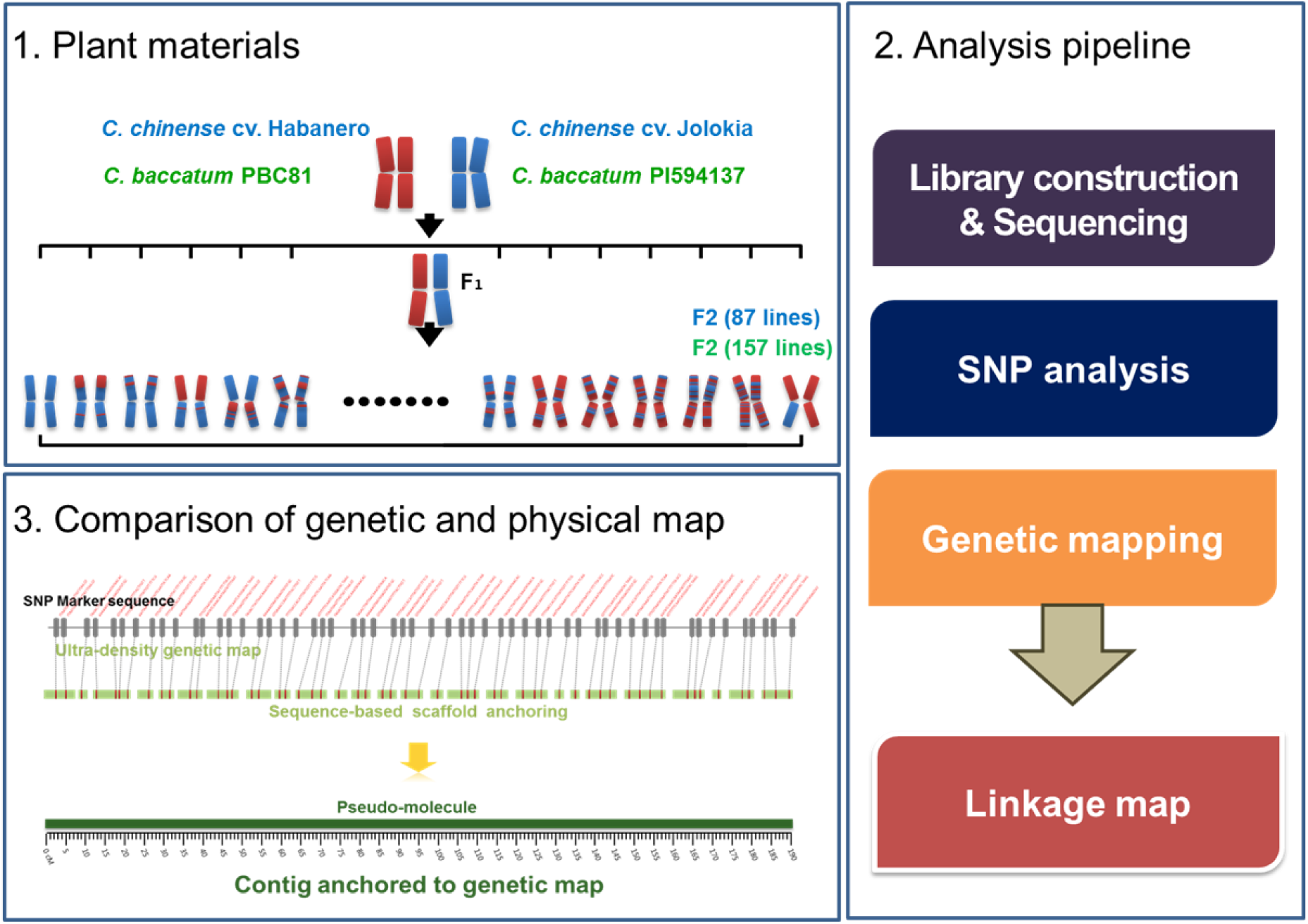
Construction of genetic maps and pseudomolecule generation for the pepper genomes. The plant materials used for construction of genetic maps of Baccatum and Chinense are shown in the first diagram. The second diagram depicts the process of genetic map construction for the two pepper genomes. The third diagram depicts the scaffold anchoring using the genetic maps for the construction of pseudo-chromosomes for the two pepper genomes.

Repeat annotation was performed with the assembled genomes and the initial contigs covering the estimated genome sizes of the three species (Extended Data Fig. 5 and Supplementary Tables 7-8). Overall, ~85% of the initial contigs were annotated as repeat sequences. LTR-Rs of the Ty3-*gypsy* superfamily accounted for about half of the entire genome in each of the three species (Supplementary Tables 7-8). Among the subgroups of the *gypsy* superfamily, *del* elements comprised the largest fraction, representing 41.5, 34.9 and 41.7% (1,482, 1,337 and 1,343 Mb) in Annuum, Baccatum and Chinense, respectively. Furthermore, *athila* elements were more abundant (>2 fold) in Baccatum, indicating that the *athila* subgroup contributed to species-specific genome expansion in the Baccatum lineage (Supplementary Table 8).

**Extended Data Figure 5.**
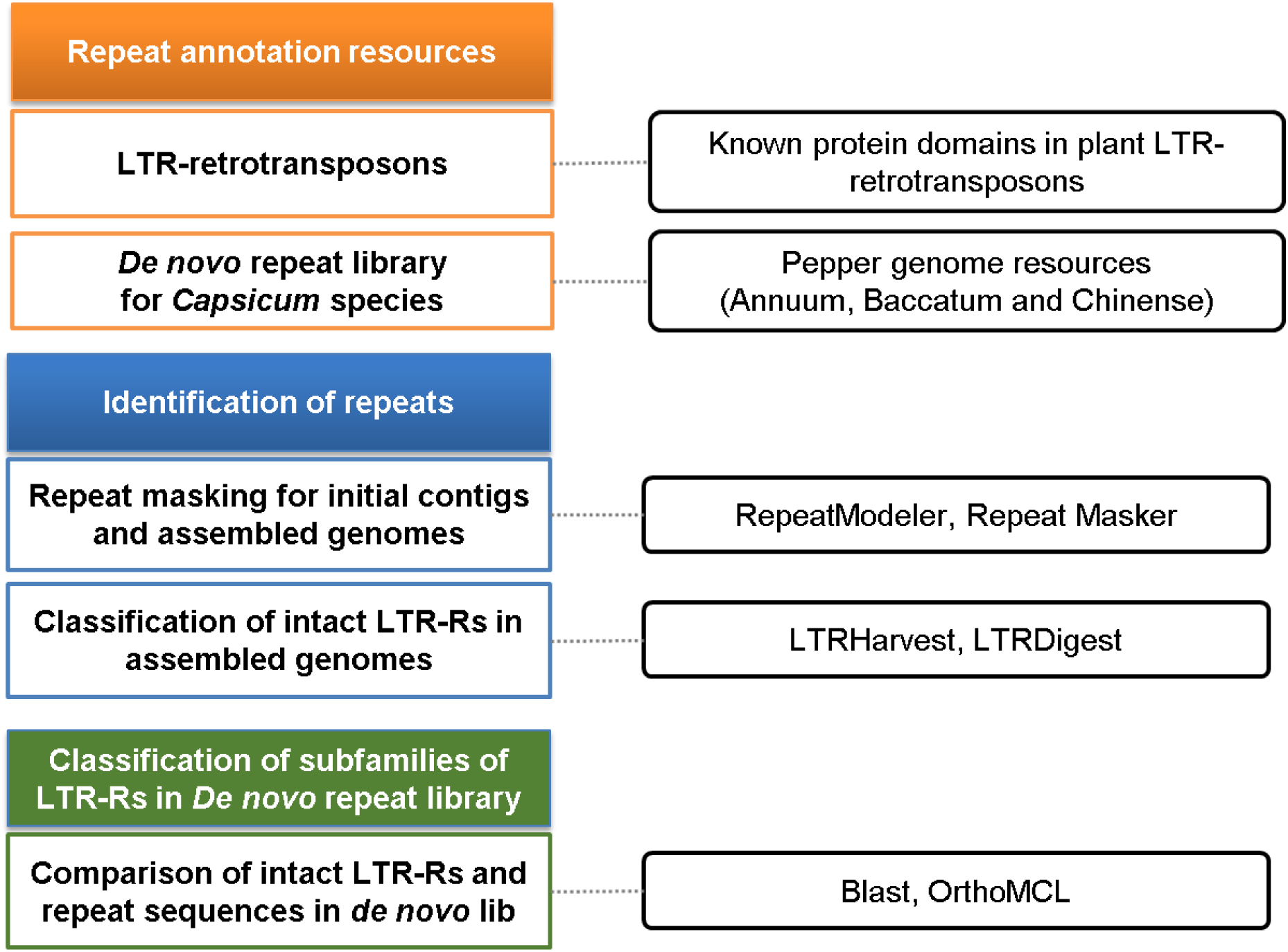
The pipeline used for annotation of repeats in the pepper genomes. The diagram depicts three steps in repeat annotation, i) construction of repeat libraries using repeat annotation resources, ii) identification of repeats and intact LTR-Rs in the pepper genomes, and iii) assignment of family information for LTR-Rs in the *de novo* repeat library. The diagrams in the right panel indicate the resources and tools used for the annotation of repeats (See the Methods).

### Speciation and evolution of the *Capsicum* species

A phylogenetic analysis of the peppers with other plant species revealed that the divergence among the three peppers occurred first between Baccatum and a progenitor of the other two peppers ~1.7 million years ago (MYA), followed by divergence between Annuum and Chinense lineages ~1.1 MYA (Fig. 1a and Supplementary Table 9). To identify genomic changes in the three pepper species, we compared the genome structures, LTR-R insertion patterns, and gene duplication histories across these pepper genomes (Figs. 1b-c and 2).

**Figure 1.**
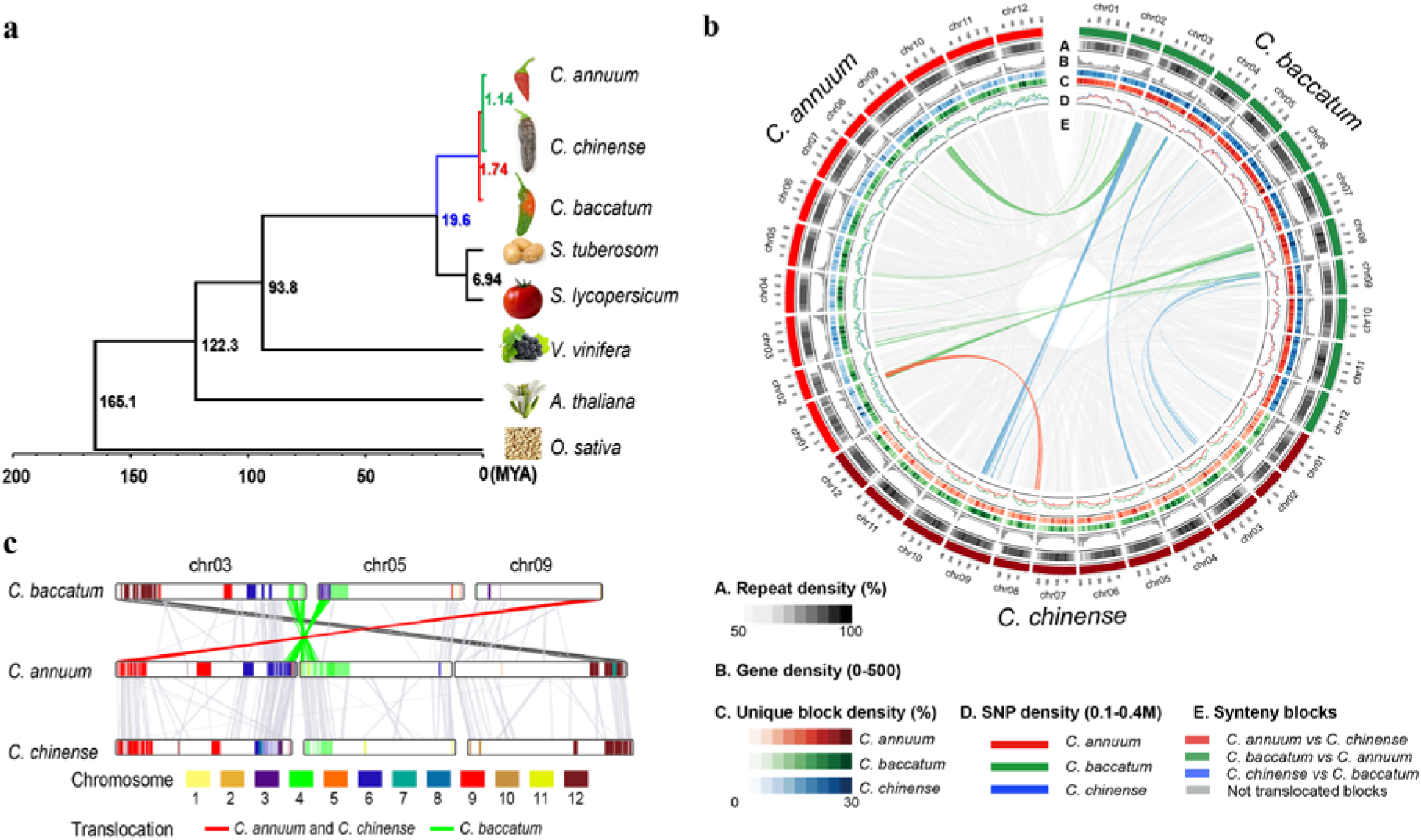
Lineage-divergence and genome structure comparisons of three *Capsicum* species. **a,** The reconstructed phylogenetic tree of eight plant genomes indicates their evolutionary relationships and estimated divergence times. **b,** The circular diagram shows the distribution of repeats, genes, genomic variations, and genome rearrangements in the pepper genomes. The subcategories indicate the density of repeats (A), genes (B), species-specific blocks (C), and SNPs (D) in the pepper genomes. The subcategory E depicts collinear and translocated blocks among the pepper genomes. **c,** A linear comparison of the rearranged blocks in the pepper genomes. Colours in the bars indicate translocated regions when comparing to tomato and potato genomes. The line colours indicate translocations in the ancestral lineage leading to Annuum and Chinense (red), in Baccatum (green) and in the ancestor of Annuum and Chinense or Baccatum (dark grey).

**Figure 2.**
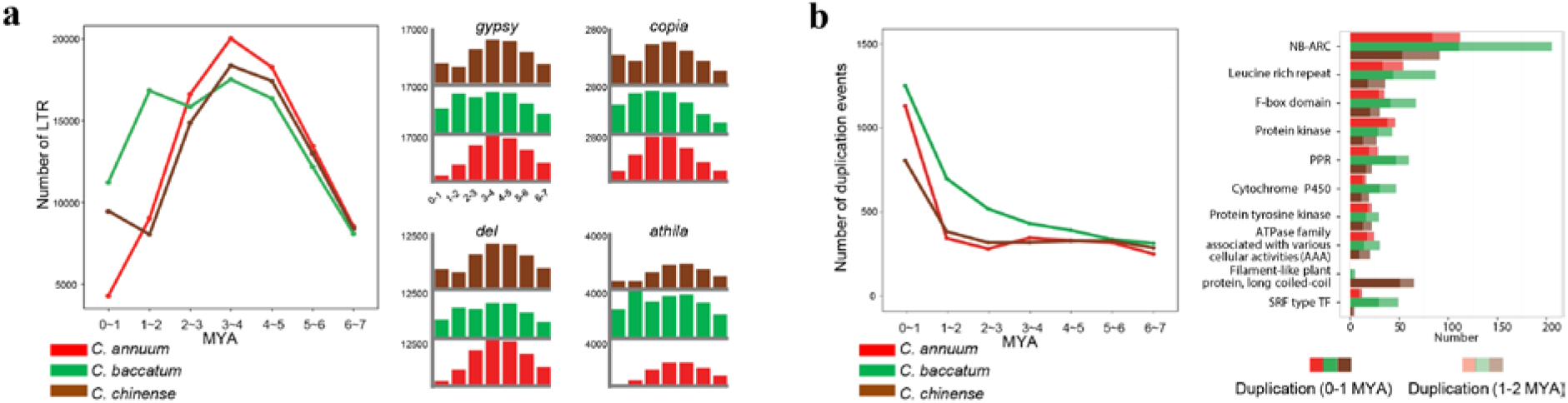
Evolutionary history of LTR-Rs and duplications of protein-coding genes in the pepper genomes. **a,** Distribution of LTR-R insertions. The graphs in the left and right panels depict the predicted insertion dates of LTR-R superfamilies (*gypsy, copia*) and two specific families (*del, athila*). The *x-* and *y*-axes indicate the insertion times and the number of insertions at each time, respectively. **b,** Time-scaled gene duplication history (left panel) and top 10 repertoires of massive gene duplication (right panel). The *x-* and *y*-axes of the graph in the left panel indicate the approximate duplication time (MYA) and the number of gene duplications, respectively. The *x-* and *y*-axes of the histogram in the right panel represent the number of genes and domain description, respectively.

Chromosomal rearrangement is an important force in speciation, often producing unbalanced gametes that reduce hybrid fertility^28^. We performed an inter-genomic structural comparison and detected translocations among the pepper genomes (Fig. 1b). The results show that chromosomes 3, 5 and 9 exhibit translocations that differentiate Baccatum from the other two species (Fig. 1b-c). Collinearity comparisons among *Capsicum* species and two *Solanum* species revealed that the distal region on the long arm of chromosome 9 was conserved in Baccatum but translocated to the short arm of chromosome 3 in a shared ancestor of Annuum and Chinense (Fig. 1c and Extended Data Fig. 6). Furthermore, chromosomes 6 and 4 of *Solanum* were detected in the terminal regions of the long and short arms of chromosomes 3 and 5 in Annuum and Chinense, respectively. In contrast, the orthologous regions of *Solanum* were mixed in the corresponding blocks of Baccatum (Fig. 1c and Extended Data Fig. 6). This indicates that the distal regions of the long and short arms of chromosomes 3 and 5 were translocated in the Baccatum lineage. We detected translocations between the terminal regions of the short arm of chromosome 3 in Baccatum and the long arm of chromosome 9 in Annuum and Chinense. Consequently, our analyses revealed that translocations have generated hetero karyotypes in both the Baccatum and the Annuum/Chinense progenitor lineages.

**Extended Data Figure 6.**
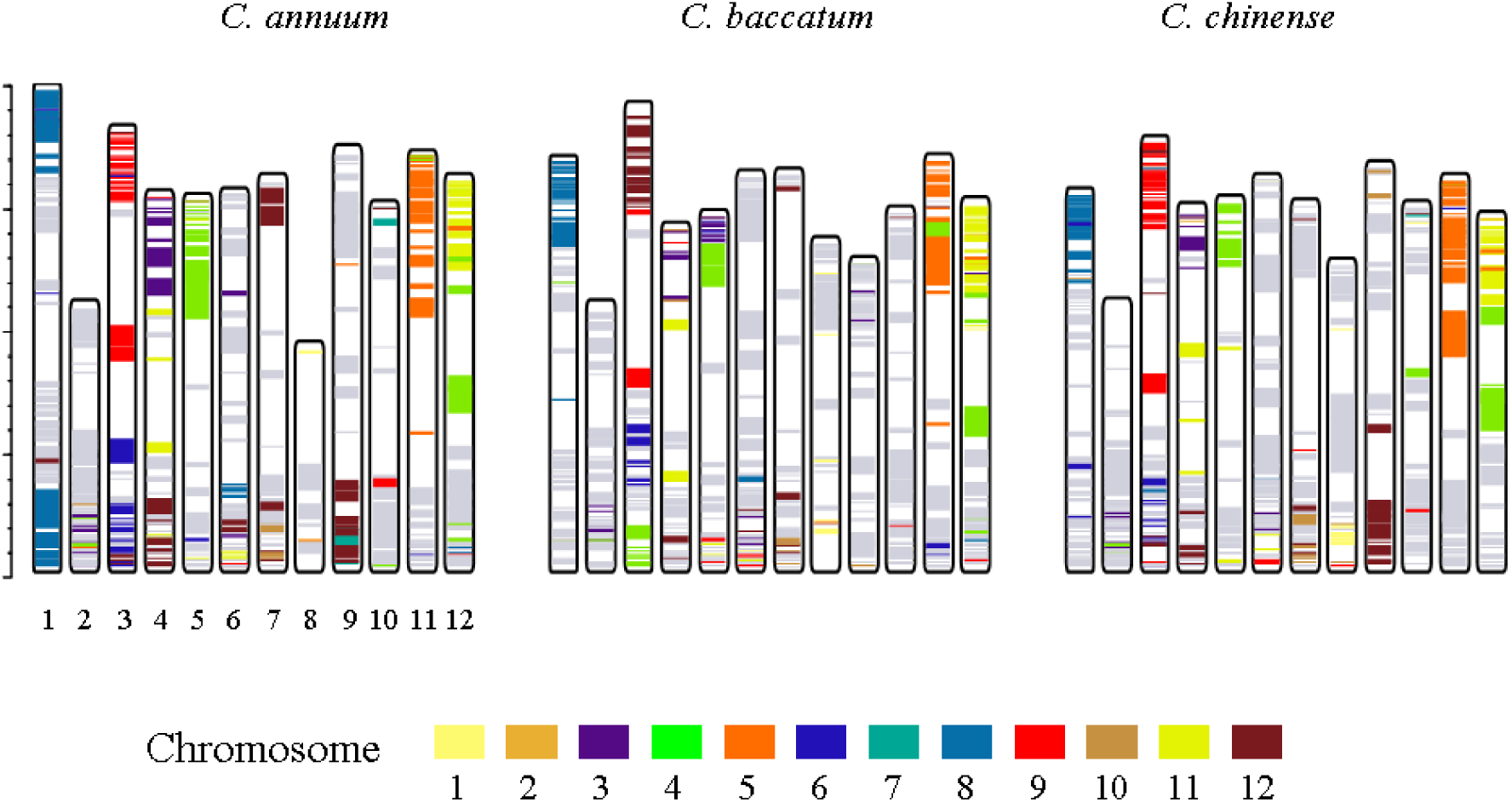
The collinear blocks in pepper genomes exhibiting conserved syntenic regions between tomato and potato genomes. The colours in the bars indicate collinear blocks in the pepper genomes that are also conserved as blocks within the tomato and potato genomes. Chromatic and grey clolurs indicate translocated and non-translocated regions, respectively, between the *Capsicum* and *Solanum* genomes.

To compare LTR-R insertion patterns across the pepper genomes, we identified full length LTR-Rs in each assembled genome and calculated their insertion times^29^ (Extended Data Fig. 5 and Supplementary Table 10). A peak of LTR-R activity in Baccatum appeared around its speciation time 1 to 2 MYA (Fig. 2a). Especially, the *athila* family was highly amplified in Baccatum around the estimated speciation time, indicating that this subgroup may have explosively increased in Baccatum after speciation. In Chinense, we observed the recent proliferation of LTR-retrotransposons (Fig. 2a).

Gene duplication is a major mechanism generating functional diversity between species by the creation of new genes^30,31^. We detected recent gene duplication events and characterised the repertories of duplicated genes in the three pepper genomes during and after speciation (Fig. 2b). Overall, the duplication events were particularly frequent in the Baccatum lineage, both during and after the speciation. In particular, NLRs were extensively amplified in Baccatum in the last 0-2 MYA, and more recently in the other two peppers (Fig. 2b). Taken together, those results suggest that the chromosomal rearrangements, accumulation of specific LTR-Rs, and differential gene duplications have contributed enormously to genome diversification in the *Capsicum.*

### Massive creation of new NLRs via LTR-R-driven retroduplication

Previous study suggested that NLRs were amplified in pepper compared to tomato and potato genomes^21^. In particular, coiled-coil NLR subgroup 2 (CNL-G2) was highly expanded in the pepper genome (Supplementary Table 11). To explore the possible mechanism of the NLR proliferation in *Capsicum* spp., we analyzed the NLRs and their flanking sequences. We identified 157, 163 and 116 NLRs located inside LTR-Rs in Annuum, Baccatum and Chinense, respectively (Supplementary Tables 11-13). Hence, a large proportion (~18%) of the NLRs were amplified by LTR-Rs, with the structures indicating that their retroduplicated origin is still intact. Most of these NLRs (~71%) were in the CNL-G2 category, indicating that the copy number of specific NLR subfamilies was particularly expanded (Fig. 3a). Furthermore, most of the retroduplicated NLRs (~71% of the total and ~66% of the CNL-G2 type) were inside non-autonomous LTR-Rs that contained no *gag* or *pol* protein coding potential (Supplementary Table 14). This suggests that all steps for the retroduplication, presumably including the initial sequence capture process, had to be provided *in trans.* When we compared the retroduplicated and other NLRs in the CNL-G2 category, the number and length of exons were significantly fewer and longer in the retroduplicated NLRs, but not all of these were single-exon genes (Fig. 3b-c). In total, ~44% of the retroduplicated NLRs in each species had multiple exons but all of those had a reduced number of introns compared with their predicted parental sequences (Supplementary Tables 13 and 15).

**Figure 3.**
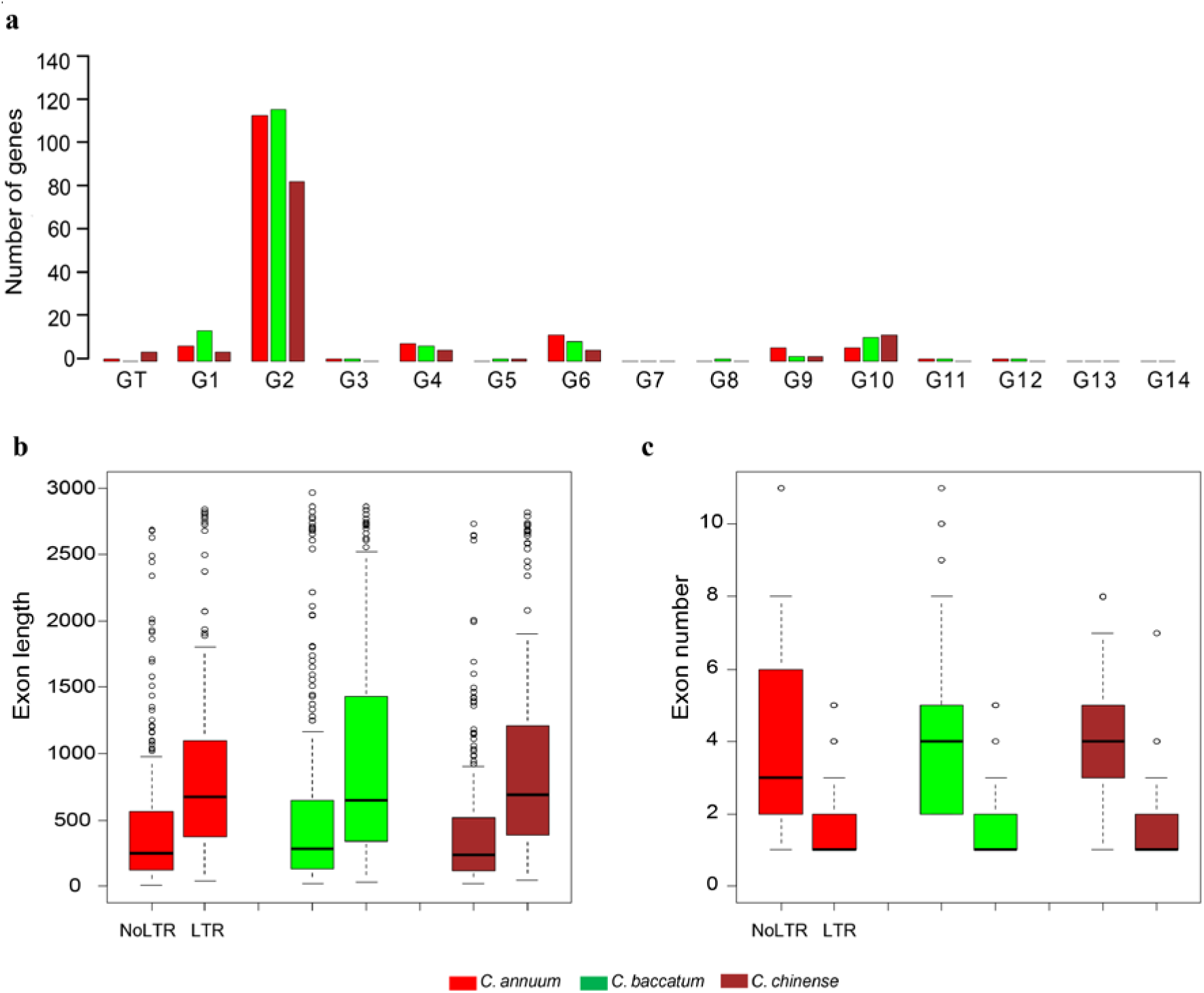
Emergence of large NLR gene families by retroduplication. The bar graph indicates the number of retroduplicated NLRs in each subgroup. **a,** The bar graph indicates the number of retroduplicated NLRs in each subgroup. The *x-* and *y*-axes indicate subgroups and the numbers of genes, respectively. **b-c,** The exon lengths and the numbers of normal and retroduplicated NLRs are depicted. **b,** The *x-* and *y*-axes indicate the normal and retroduplicated NLR groups and their exon lengths, respectively. **c,** The *x-* and *y*-axes mean the groups of NLRs and the exon numbers, respectively.

Earlier analyses of the tomato, potato and rice genomes indicated that retroduplication is a general feature of genome evolution in the plant kingdom (Supplementary Table 11). We found that 22, 103 and 30 (8, 23, and 6%) of NLRs were inside LTR-Rs in tomato, potato and rice, respectively. Of these, we identified parental sequences with multiple exons for 18, 88 and 22 of the NLRs inside LTR-Rs in tomato, potato and rice, respectively, thus confirming their origin by retroduplication (Supplementary Table 16). Similar to the peppers, NLRs in particular subgroups were primarily retroduplicated in potato but the duplicated subgroups were distinct in each species (Supplementary Table 11). These results indicate that LTR-Rs played a key role in the expansion of NLRs by retroduplication throughout the plant kingdom, and that the detected events are both recent and lineage-specific.

In addition to the NLRs, we looked for other genes inside LTR-Rs in the six plant species (Supplementary Table 17). In total, a range from 1,398 genes in rice to 3,898 genes in potato genomes were found to be inside LTR-Rs, suggesting that 4 to 10% of all the genes in these plant species emerged by LTR-R-driven retroduplication. On average, ~45% of them had functional domains including highly amplified families such as MADS-box TFs, cytochrome P450s, and protein kinases, and ~42% of those genes were expressed in one or more investigated tissues by RNA-Seq analysis (Supplementary Table 17).

### Evolutionary mechanisms for the emergence of disease resistance genes in Solanaceae

The *L* genes encoded by the NLRs are known to render resistance in peppers against *Tobamoviruses* and they belong to the CNL-G4 category, along with *I2* in tomato that provides resistance to race 2 of *Fusarium oxysporum* f. sp. *lycopersici* and *R3a* in potato that provides resistance to the late blight pathogen, *Phytophthora infestans*^32-34^. Each of those is a single exon gene encoding a peptide of ~1,300 amino acids. Synteny analysis and sequence comparison among pepper, potato and tomato genomes suggested *L*, *I2*, and *R3a* are orthologous genes and the genomic regions containing *L*, *I2* and *R3a* were tightly linked (Extended Data Fig. 7a and Supplementary Table 18). These results suggest the possibility that the genes originated by an early retroduplication, and then underwent divergent evolution in each lineage.

**Extended Data Figure 7.**
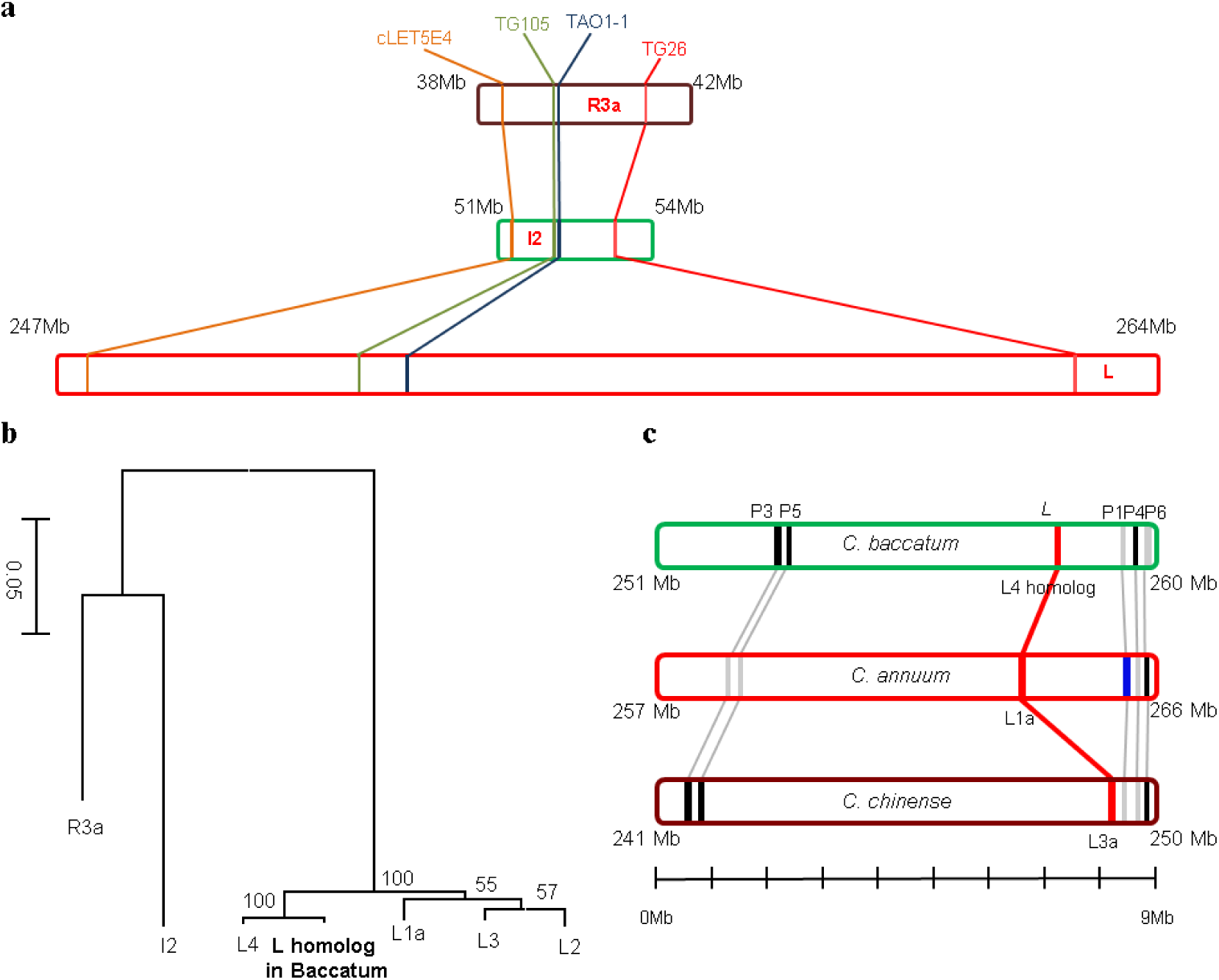
Comparative analyses of *R3a, I2* and *L* genes and the locations of those genes in the Solanaceae genomes. **a,** Syntenic regions including *L*, *I2*, and *R3a* in the pepper, tomato and potato genomes are depicted as bar graphs. The marker names were described in a previous study^33^. **b,** The phylogenetic tree of *L*, *I2* and *R3a.* **c,** Bar graphs depict the locations of the closet homologs (P1 to P6) of the *L* genes in the pepper genomes as candidate parental sequences. Black and grey lines in the bars mean the presence and absence of the parental sequences in pepper genomes, respectively. The blue band within the red bar indicates the location of P1 in Annuum, the most likely parental gene of *L1a.*

We examined the evolutionary history of *L* genes with their parents in the pepper genomes (Fig 4, Extended Data Fig. 7b-c, and Supplementary Tables 19-20). The candidates of a parental gene (P1 to P6) were identified considering similarity, *Ks* values (synonymous substitutions/synonymous site), and alignment coverage to *L* genes. All candidate parental sequences contained multiple exons. When candidate parental sequence P1 was compared with *L* in Annuum, the results suggested that *L* was derived from retroduplication in the ancestral lineage of *Capsicum* spp. ~8.9 MYA (Fig. 4). Because *L* has 6.7 kb coding exons, with only an intron in the 3’ UTR, and the presence of both flanking direct repeat sequences and a poly(A) “tail” encoded in the DNA, our analysis suggests that *L* emerged through capture and reverse transcription by a long interspersed nuclear element (LINE)-driven retroduplication (Fig. 4 and Extended Data Fig. 8). Sequence comparison of *L* genes in the three genomes and *L4* in *C. chacoense* revealed that the *L* genes were diversified by accumulation of lineage-specific sequence mutations after speciation within *Capsicum* (Fig. 4 and Supplementary Table 21). Consequently, our results suggest that the ancestor of the *L* genes was derived from retroduplication and that subsequent divergent evolution has led to race-specific resistance against diverse strains of *Tobamovirus* in each species of *Capsicum* after speciation (Fig. 4).

**Figure 4.**
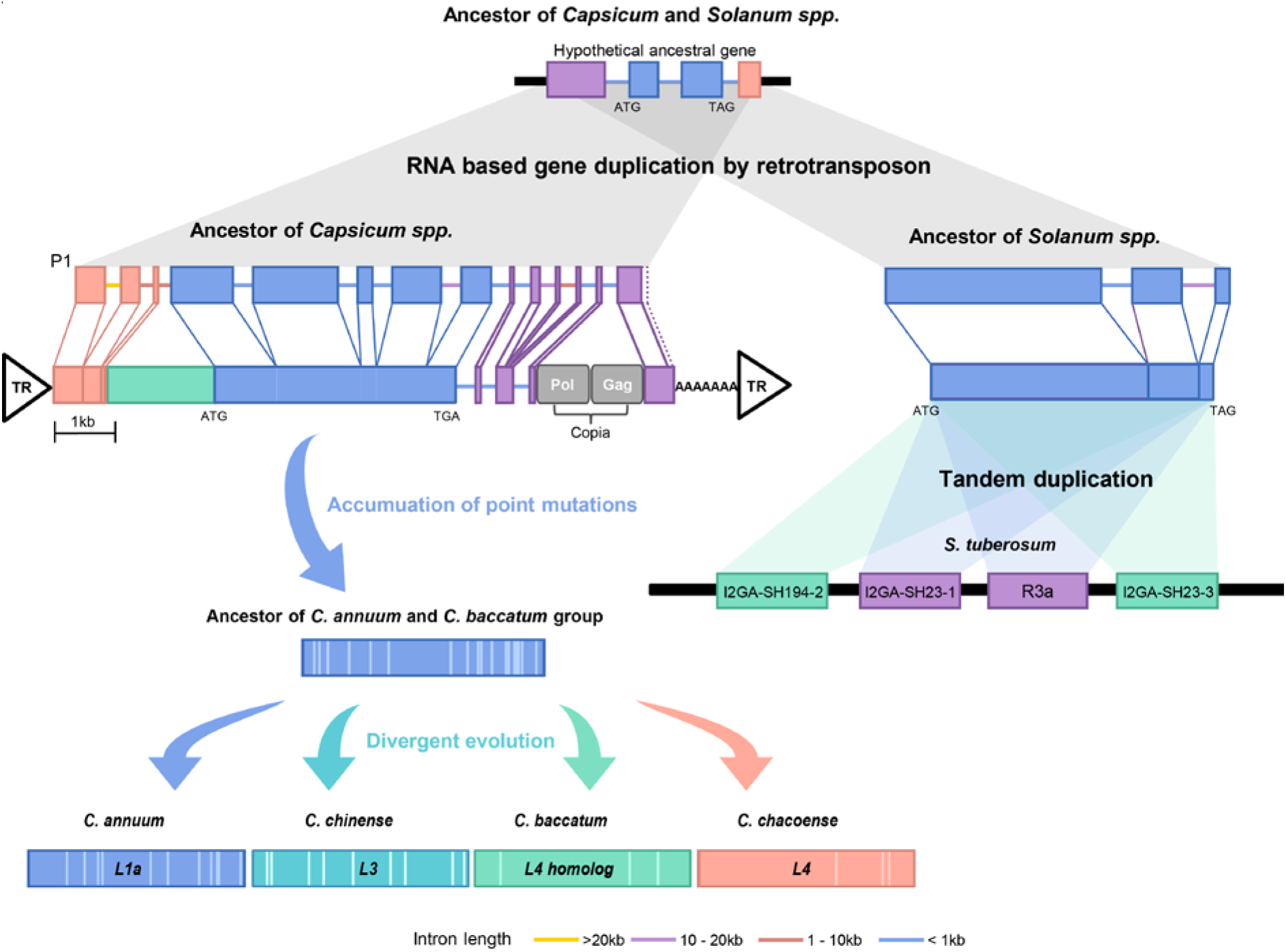
Emergence and evolution of *L* and *R3a* genes in the pepper and potato genomes. Models for the evolution of *L* and *R3a* in the pepper and potato are depicted. The gene names in the *R3a* cluster are from the previous analysis of Huang *et al*^33^. The model proposes that *L* and *R3a* gene ancestors were first created by retroduplication, followed by the accumulation of point mutations and tandem duplication, respectively. DNA sequence indicative of a poly(A) tail and flanking terminal repeat (TR) sequences are depicted in the diagram as genomic evidence for a retroduplicated origin of *L*.

**Extended Data Figure 8.**
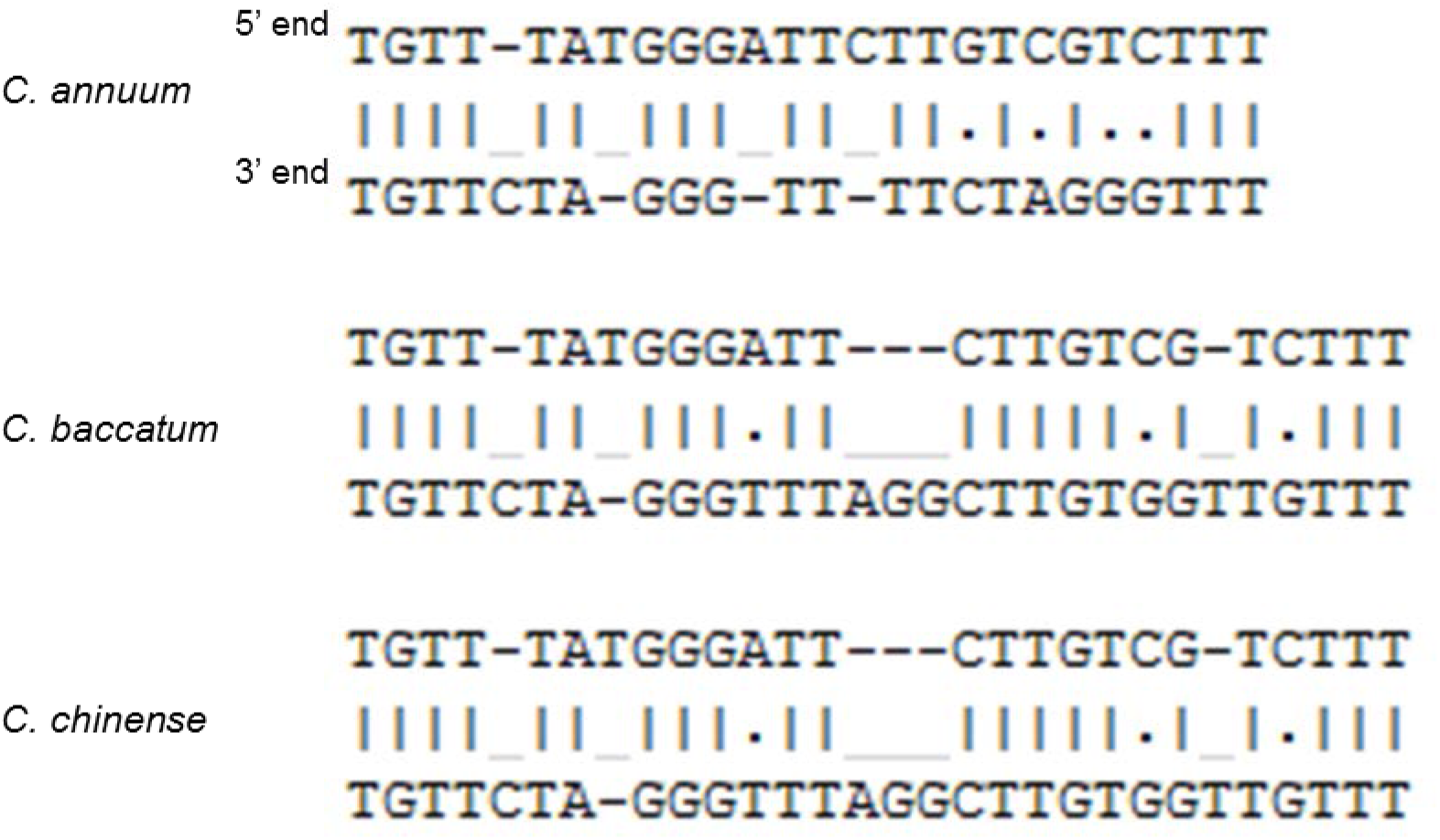
Alignments between the 5’ and 3’ end direct repeat sequences flanking the *L* genes in the pepper genomes.

To analyse the evolutionary processes acting on *R3a* of potato, we first performed a genome-wide search for the *R3a* as well as for candidate parental sequences. Because *R3a* is absent in the current potato reference genome^35^, we could not carry out accurate comparisons of *R3a* and their homologues. However, *R3a* and its clustered genes originated from wild species, *Solanum demissum^36^* were available in a public database. So, we compared these sequences with their closest homologs in the reference potato genome^33^. Our analyses revealed that intronless sequences of the ancestral potato *R3a* might have emerged by RNA-based gene duplication in a shared ancestor of potato and tomato (Fig. 4). Subsequently, *R3a* and its paralogues were amplified by two rounds of tandem gene duplication after the divergence of potato and tomato (Fig. 4 and Supplementary Table 22). Taken together, our results suggest that retroduplication events are a main evolutionary process in the emergence of new plant disease resistance genes, which can gain function via subsequent sequence variation and tandem duplication.

### Evolution of potential anthracnose resistance genes in Baccatum

Pepper anthracnose caused by *Colletotrichum* spp. is one of the most devastating diseases in worldwide pepper production^37^. Due to the complexity of the interactions between the host and *Colletotrichum* spp. and the lack of resistance in the Annuum gene pool, a few Baccatum varieties were identified as the only breeding resources for anthracnose resistance^38^. Using pre-existing genetic information^39^, we identified the pertinent genomic regions and obtained 64 NLRs from a 3.8 Mb region of Baccatum chromosome 3 as candidate resistance genes for *C. capsici* (Fig. 5 and Supplementary Table 23). Previous studies reported that the main quantitative trait locus (QTL) for pepper resistance against *C. capsici* was located on chromosome 9^39^, however, we found that QTL is located on chromosome 3 due to translocation in Baccatum and Annuum (Fig. 1c). We obtained 35 Baccatum-specific NLRs (27 in CNL-G2, 5 in CNL-G10 and 3 in CNL-G10) from the 64 NLRs by sequence comparison among the three pepper genomes (Fig. 5). Considering the gene duplication history, 15 of the 35 genes appear to have emerged after generation of the Baccatum lineage and all of them belong to the CNL-G2 category. Transcriptome evidence indicated that 10 of those 15 genes are expressed in one or more tissues (Fig. 5 and Supplementary Table 23). Furthermore, half of the 15 genes appear to have emerged by retroduplication (Fig. 5). Consequently, our results suggest that retroduplication along with tandem and segmental duplications, has played a major role in the emergence of anthracnose resistance genes in the Baccatum lineage.

**Figure 5.**
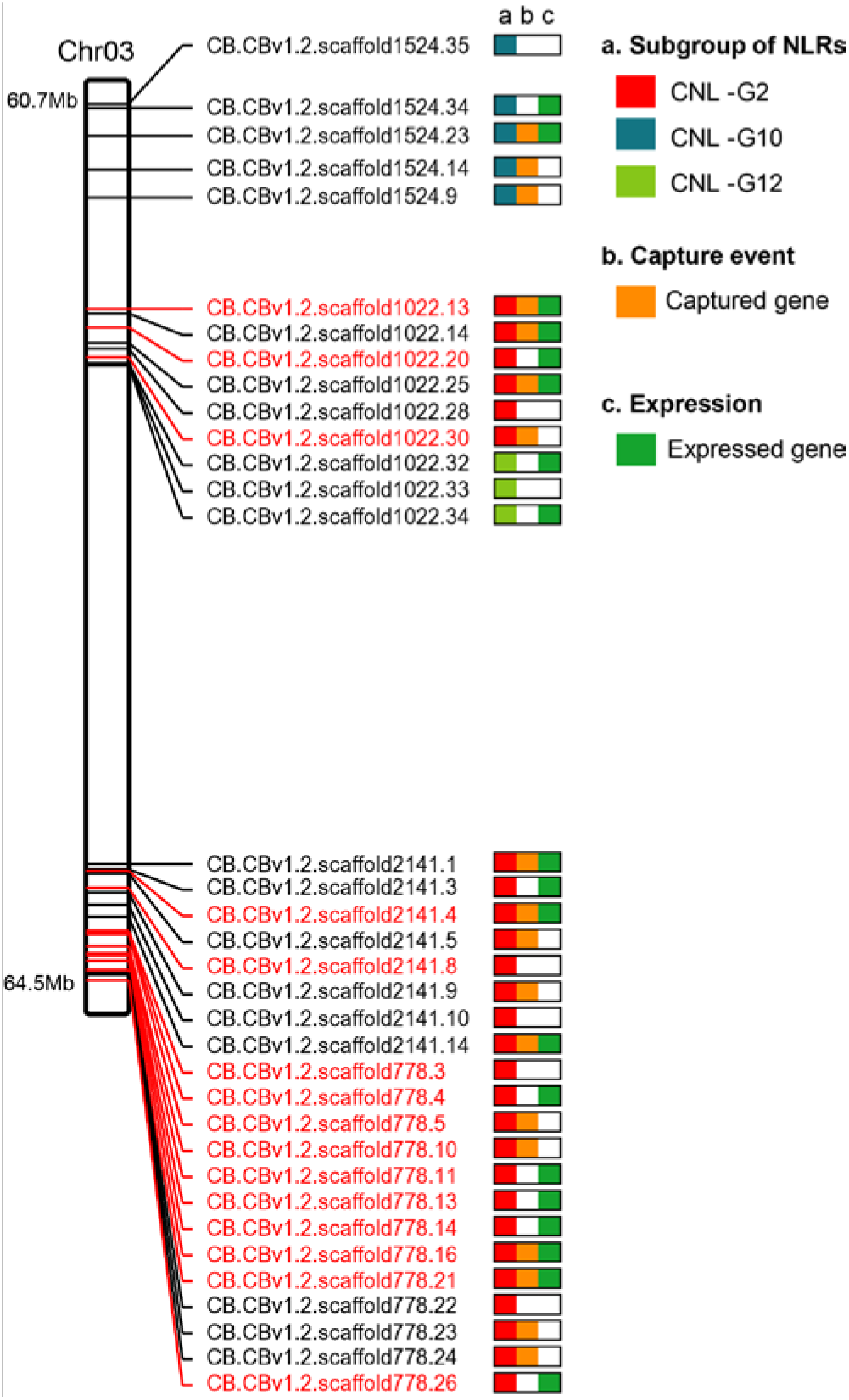
Potential *C. capsici* anthracnose resistance genes for in chromosome 3 of Baccatum. Baccatum-specific NLRs in the major QTL region are visualised on 3.8 Mb of chromosome 3. The chromosome plot shows the subgroups, proposed retroduplication events, and expression results for the NLRs. The black and red texts indicate NLR IDs that emerged before and after the speciation of Baccatum, respectively.

## Conclusions

In this study, we generated new and improved genome resources for three *Capsicum* species. Our data provide an accurate and updated gene model of the pre-existing reference pepper genome based on annotation with accumulated knowledge, highlighting the importance of genome improvement after the completion of sequencing project. High-quality pseudo-chromosomes constructed from three pepper genomes enabled precise comparisons of genome structures and evolutionary analyses, providing new insights into interspecies diversification via genome rearrangements and lineage-specific evolution of LTR-Rs and genes in the genus *Capsicum.* Furthermore, we found evidences of the massive evolution of NLRs by LTR-mediated retroduplication in dicot Solanaceae and monocot rice, suggesting that such phenomena are ubiquitous in angiosperms. Our results suggested that at least 6 to 23% of plant NLRs were emerged by LTR-R-driven duplication (Supplementary Table 11). A notable feature of this retroduplication is that distinct subfamilies of NLRs were highly retroduplicated in different plant lineages. We unveiled the emergence of functional disease resistance genes in the Solanaceae family by retroduplication and the subsequent neofunctionalisation of those genes by dynamic evolutionary processes including lineage-specific sequence mutation and tandem duplication. Our data suggest that a large proportion of all genes (~4 to 10%) in plant species might have emerged by LTR-R-driven retroduplication. We observed the lineage-specific amplification of specific gene families by LTR-Rs in various plant species, including such genes as those encoding cytochrome P450s in potato and MADS-box TFs in Baccatum (Supplementary Table 17). Taken together, our results provide new insights into the evolution of functional plant disease-resistance genes that belong to the NLR family as well as other high copy number gene families that are present in the plants.

### Online Content

The Methods and Supplementary Information with any associated references are available in the online version of the paper.

## URLs

GapCloser v1.12, http://soap.genomics.org.cn/down/GapCloser_release_2011.tar.gz

RepeatModeler, http://www.repeatmasker.org/RepeatModeler.html

Rice gene expression data, http://rice.plantbiology.msu.edu/

Potato gene expression data, http://solanaceae.plantbiology.msu.edu/pgsc_download.shtml

## Acknowledgements

This work was supported by a grant from the Agricultural Genome Center of the Next Generation Biogreen 21 Program of RDA (Project No. PJ01127501) and by a grant from the Ministry of Science, ICT, and Future Planning (MSIP) of the Korean government through the National Research Foundation (NRF-2015R1A2A1A01002327) to D.C.

## Author contributions

D.C. conceived the project, designed the content, and organised the manuscript. S.K. performed data generation/analysis and managed subprojects. S.-B.K., H.-A.L., and H.-Y.L. prepared the DNA and RNA samples. S.K., K.C., K.H., and J.L. performed the *de novo* genome construction of two domesticated pepper genomes and the improvement of pre-existing reference genome. S.K., M.-S.K., J.P., Y.-M.K., N.K., and R.W.K. carried out the gene annotation. S.K., J.-K.L., S.-C.L., T.-J.Y., J.K., H.-Y.L., and B.-C.K. fulfilled the repeat annotation and analyses. S.K., H.-S.S., J.J., J.H.H., and Y.-H.L. implemented the genome structure comparison and evolutionary analyses of TEs and genes. S.K., E.S., J.P., Y.-S.L, S.O., J.H.L., E.C., E.C., S.E.L., G.C., H. S., and W.-H.K. performed the gene family analyses. S.K., E.S., and J.P. carried out the retroduplication analyses for the NLRs and other gene families. S.K., H.K., H.-S.S., and Y.H. designed and visualized the figures. S.K., S.-I.Y., Y.-M.K., K.-T.K., J.-M.L., M.P., J.L.B., and D.C. wrote the manuscript.

## Author information

The genome sequences of *C. chinense* and *C. baccatum* are deposited in the GenBank under the accessions MCIT00000000 and MLFT00000000 (the versions described in this paper are version MCIT01000000 and MLFT01000000). Further information, containing pseudomolecules and annotations is uploaded to our website (http://peppergenome.snu.ac.kr). The authors declare no competing financial interests. Correspondence and requests for materials should be addressed to D.C.

## Methods

### Genome assembly

In total, 425.7 Gb and 526.7 Gb of the Chinense and Baccatum genome sequences were generated in the Illumina Hiseq 2500 system (Supplementary Table 1). To remove unnecessary sequences for genome assembly, preprocessing analysis was performed as previously described^25^ (Supplementary Table 2). The *de novo* genome assembly of each species was performed with SOAPdenovo2^40^ using the filtered raw sequences with parameters K=77 and K=81 for Chinense and Baccatum, respectively (Supplementary Table 3). The SSPACE software^41^ was employed for additional scaffolding (-x 0 -m 46 -k 10 -a 0.4 -p 1), and Gapcloser v.1.12 (See URLs) and Platanus^42^ were implemented using default parameters to close gaps.

### Gene and repeat annotations

Gene annotation was performed for the three pepper genomes as described in Extended Data Figure 2. To annotate protein coding genes, we assembled transcripts using Tophat and Cufflinks^43^ with the RNA-Seq reads described in Supplementary Table 12 and in a previous study^25^. The ISGAP pipeline^44^ was used to extract accurate coding sequences from the assembled transcripts. Plant refSeq^45^ and the public protein databases for *Arabidopsis* (TAIR 10), tomato (iTAG 2.3), potato (PGSC v3.4), and pepper (PGA v1.55) were used with Exonerate v2.2.0^46^ to align protein to the pepper genomes. *Ab initio* prediction was carried out with AUGUSTUS^47^ version 3.0 using an in-house training set consisting of full-length cDNA generated from transcriptome analysis and by protein alignment. Consensus gene models were determined with EVM^48^ and the biological description of each gene model was assigned based on the Uniprot database and INTERPRO scan v5.15-54.0^49^.

Repeat sequences were annotated in the initial contigs representing the whole genome sizes and the assembled genomes of the three peppers, as shown in Extended Data Figure 5. An integrated repeat library of the three peppers was constructed using RepeatModeler (See URLs). Annotation of intact LTR-Rs was performed using LTRHarvest^50^ (-maxlenltr 2000 and -similar 80) and LTRDigest^51^. The subgroup of LTR-Rs in the integrated library was classified by comparing their sequences to those of the intact LTR-Rs using BLASTN (similarity >90%) (Extended Data Fig. 5).

### Comparison of genome structures

To identify regions that were either conserved or translocated between *Capsicum* and *Solanum* species, we performed collinear analysis with MCScanX^52^ using the gene models of the three peppers, and the tomato and potato genomes described in Supplementary Table 9. We identified regions that were not translocated between the tomato and potato genomes as conserved blocks in the *Solanum* species. The conserved blocks in the *Solanum* species were then compared to the three pepper genomes. Blocks in the pepper genomes that were conserved or translocated between the *Capsicum* and *Solanum* species were determined as shown in Extended Data Figure 6. To investigate the translocated blocks in the three pepper genomes, we examined the gene collinearity for syntenic blocks as shown in Extended Data Figure 6 and Figure 1c.

### Gene duplication history

To estimate the gene duplication times of the annotated genes in the pepper genomes, we constructed a computational pipeline by modifying a previously described method^53^. We first performed gene clustering analysis using OrthoMCL^54^ to classify the gene family. We assumed that the genes in the same clusters were in the same family and performed all-by-all alignments of the coding sequences within the clusters in each species using PRANK^55^. For each alignment result, the *Ks* values were calculated using KaKs Calculator,^56^ and single-linkage clustering for the *Ks* values was performed using the hclust function in the R package. The molecular clock rate (*r*) was calculated to be 6.96 × 10^-9^ substitutions per synonymous site per year^57^. The duplication time was estimated using the formula, *Ks* value / 2r.

### Estimation of divergence time

To estimate the divergence times of the plant genomes, we identified 2,540 single copy genes in the rice, *Arabidopsis* (TAIR10), grape (VvGDB v2.0), tomato (v2.3), and potato (PGSC v3.4) genomes and the three pepper genomes using OrthoMCL clustering^54^ (Supplementary Table 9). Multiple alignments of the single copy genes from the eight genomes were implemented using PRANK^55^ (-f=nexus -codon). The speciation times of the eight plant species were calculated by phylogenetic analysis using the BEAST package^58^.

### Evolutionary analyses of LTR-Rs

For the intact LTR-Rs, we performed alignment of the sequences between the 5’ and 3’ LTRs using PRANK. The DNA substitution rates (K) between the 5’ and 3’ LTRs were calculated using baseml in the PAML package^59^. The insertion times of the LTR-Rs were estimated using the formula, *K/2r* (*r* =1.3 × 10^−8^)^60^.

### Identification of retroduplicated NLRs in the plant genomes

To identify NLR genes inside LTR-Rs, we used the rice (MSU RGAP 7), potato (PGSC v3.4), and tomato (v2.3) genomes with the three pepper genomes. We first identified NLRs using a previously constructed pipeline^21^ and extracted the NLRs within putative LTR-Rs predicted by LTRHarvest (Supplementary Table 11). We then compared those results with the repeat annotated by RepeatMasker, and if the NLRs inside LTR-Rs overlapped with other TEs such as DNA transposons or Helitrons, we considered that the LTR-R predicted using LTRHarvest was probably incorrect and was then removed. Because of rapid deletion of LTR-Rs and other unselected DNA in all flowering plants^1,14^, we performed an additional identification of NLRs inside LTR-Rs using the annotated repeats including the partial LTR-Rs generated by RepeatMasker. We reasoned that if the NLRs were fully contained within LTR-Rs annotated by RepeatMasker, the NLRs were retroduplicated. To verify intron removal from the retroduplicated NLRs, we determined whether the candidate parental sequences of the NLRs contained multiple exons by aligning the candidate parental sequences with the NLRs using Exonerate^46^, requiring >95% query coverage of the NLRs (Supplementary Table 13). To increase analysis accuracy, we ignored unclear cases where no parental sequences having multiple exons were detected for NLRs inside LTR-Rs.

To predict whole genes inside LTR-Rs in the six plant genomes, we performed genome-wide identification of possible structure of LTR-Rs using LTRHarvest, taking into account rapid sequence change between the LTRs (-similar 75%, minltrlen 100). Like annotation of the NLRs inside LTR-Rs, we extracted genes within directly repeated LTR regions as putatively retroduplicated genes. For the genes inside LTR-Rs, the number of expressed genes in one or more tissues was counted using RNA-Seq data, as described in Supplementary Table 12 and in previous analyses^25,35,61,62^ (See URLs).

### Evolutionary investigation of functional disease-resistance genes in Solanaceae genomes

The *L, I2* and *R3a* genes of pepper, tomato and potato were used to investigate evolutionary processes acting on functional disease-resistance genes in the Solanaceae plants. The *L* genes in the *Capsicum* spp. were aligned to paralogues in the pepper genomes using Exonerate^46^ and closest homologs were identified (Supplementary Tables 19-20). All of the closest homologs in each species were found to contain multiple exons and a gene that we named P1 was identified as the most likely parental sequence. By comparison of the sequence divergence between P1 and its closest homologs in the other genomes, we confirmed that P1 was Annuum-specific (Extended Data Fig. 7c). The 5’ and 3’ UTRs of *L1a* annotated based on RNA-Seq evidence were also compared to the UTRs of P1 (Fig. 4).

For *R3a* in the potato, we aligned the coding sequences of *R3a* and genes within its cluster downloaded from GenBank (AY849382, AY849383, AY849384 and AY849385) to the potato genome. Because of the absence of those genes in the potato reference genome^35^, we identified the closest homologs of *R3a* except *R3a* and its clustered genes in the potato genome. The duplication time of the *R3a* family was estimated by comparison of *R3a* and its homologs identified in the potato genome with the clustered genes. The *I2* sequence of tomato downloaded from GenBank was also used to search in tomato reference genome, but *I2* was not found.

### Identification of potential anthracnose resistance genes

To obtain candidate anthracnose resistance genes for *C. capsici*, we extracted NLRs located in the terminal region of the short arm of chromosome 3 of Baccatum based on pre-existing genetic information^39^ (Supplementary Table 23). Candidate genes that may provide resistance in Baccatum against *C. capsici* were determined based on further inspected for the sequence conservation and the gene duplication time (Fig. 5).

